# The role of parasympathetic mechanisms in the infarct-limiting effect of SGLT2 inhibitor ertugliflozin

**DOI:** 10.1101/2021.10.01.462765

**Authors:** MV Basalay, S Arjun, SM Davidson, DM Yellon

## Abstract

**Introduction:** Based on data that outcome in patients with acute myocardial infarction is predicted by final infarct size (IS), reducing IS is of paramount importance. Recent experimental studies have demonstrated a strong infarct-sparing effect of SGLT2 inhibitors – a class of drugs which have proved to be safe and beneficial in patients with heart failure. Repurposing SGLT2 inhibitors for the benefit of patients presenting with acute myocardial infarction should be preceded by investigation of the underlying mechanisms of this infarct limitation. Experimental and clinical data indicate a potential role for autonomic modulation in these mechanisms, specifically sympatho-inhibition. The aim of this study was to investigate the role of the parasympathetic mechanism in the infarct-limiting effect of SGLT2 inhibition.

**Methods:** Fortyeight Sprague Dawley male rats were fed with a standard diet containing either the SGLT2 inhibitor ertugliflozin or vehicle, for 5-7 days. Myocardial ischaemia/reperfusion injury was initiated by a 40-min occlusion of the left anterior descending coronary artery followed by a 2hr period of reperfusion under isoflurane anaesthesia. Bilateral cervical vagotomy was performed 10min prior to myocardial ischaemia. Alternatively, muscarinic receptors were blocked systemically with the non-selective blocker atropine sulphate (2 mg/kg bolus, then 1 mg/kg/h) or the M3-selective blocker 4-DAMP (2 mg/kg bolus).

**Results:** Pre-treatment with ertugliflozin reduced IS in comparison with the vehicle-treated controls (p<0.001). Bilateral vagotomy, atropine sulphate and 4-DAMP all abolished this infarct-limiting effect (IS 35±10%, 44±8%, and 35±4% respectively; P<0.01 *vs*. Ertu for vagotomy, P<0.001 *vs*. Ertu for both atropine sulphate and 4-DAMP).

**Conclusion:** These results suggest that the Infarct-limiting effect of the SGLT2 inhibitor ertugliflozin, may be mediated via activation of the vagus nerve and M3-cholinoreceptors.

## Rationale

Coronary heart disease, either in its acute or chronic form, is the leading cause of death in developed countries. The most common mechanism of acute coronary syndrome with ST-segment elevation, is obstruction of a coronary artery, resulting in severe ischaemia which rapidly develops into myocardial infarction. Coronary revascularization is necessary to reperfuse the myocardium and limit the extent of infarction, even though reperfusion causes some injury itself, which is known as reperfusion injury. Final infarct size predicts long-term clinical outcome in patients with ST-elevation myocardial infarction^1^. Therefore, finding ways to minimize such injury is vital in reducing mortality and morbidity in these patients^2^.

Inventing new drugs is a long and expensive process, while repurposing existing drugs saves costs and time for approval^3^. In this regard, two recent large-scale phase 3 placebo-controlled trials (EMPEROR-Reduced and DAPA-HF) have shown that sodium-glucose co-transporter- 2 (SGLT2) inhibitors are safe and reduce the combined risk of cardiovascular death or hospitalisation for heart failure with reduced ejection fraction, even in those without diabetes^4,5^. Based on these results, SGLT2 inhibitors are currently being widely introduced to everyday clinical practice to improve the prognosis in patients with heart failure^6^. In a recent meta-analysis of 7 pre-clinical studies in mice, rats and pigs, it was shown that, apart from improving myocardial function and prognosis in patients with heart failure, SGLT2 inhibitors can significantly reduce infarct size (IS) in regional myocardial ischaemia/reperfusion models^7^.

The mechanism of the infarct-limiting effect of this group of drugs is not clear, and is likely to be indirect, because SGLT2 is not expressed in human myocardium^8,9^. In line with this, while SGLT2 inhibitors reduce IS when administered *in vivo*, they do not protect the *ex vivo* (isolated) perfused rat heart, suggesting that the cardioprotective effect of SGLT2 inhibitors is dependent on a systemic mechanism^10^. One interesting possibility is that this systemic mechanism is neurally mediated.

If a neural mechanism is required for cardioprotection by SGLT2 it might involve the sympathetic or parasympathetic arms. Recent experimental^11,12,13^ and clinical data^14^ strongly suggest that SGLT2 expression may be up-regulated by the sympathetic nervous system^11,13^, and, *vice versa*, SGLT2 inhibition is associated with sympatho-inhibition^12,14^. However, sympatho-inhibition by SGLT2 inhibitors has only been directly demonstrated in kidneys^12^, and not in the heart^12,14^. In addition, it cannot be excluded that inhibition of sympathetic tone by this class of drugs might be secondary to improvements in haemodynamics and metabolic stress, and not causally related to cardioprotection^15^. This hypothesis is corroborated by the fact that no studies, to the best of our knowledge, have been able to demonstrate that sympathetic denervation of the heart reduces infarct size. By contrast, an increase in parasympathetic flow to the heart by electrical stimulation of the vagus nerve has been demonstrated to reduce infarct size in mice^16,17^, rats^18,19,20^, pigs^21,22^, and dogs^23^. Furthermore, activation of muscarinic acetylcholine (ACh) receptors, either their M2 or M3 subtype, is known to trigger cardioprotective signaling cascades alleviating ischaemia/reperfusion injury^24,25^. This provides a strong basis to hypothesize that cardioprotection by SGLT2 inhibitors, specifically, their infarct-limiting effect, may involve increased parasympathetic flow.

## Aim

To investigate the role of the parasympathetic nervous system in cardioprotection by SGLT2 inhibitors.

## Materials and methods

### Animals used

Sprague Dawley (SD) rats of 150-174 g weight were purchased from Charles River Laboratories, and allowed to acclimatise under standard conditions for at least 1 week.

### Drug administration

Pre-treatment with ertugliflozin (Ertu) or vehicle was performed for 5-7 days using re-pelleted rodent chow containing ertugliflozin at a dose based on the average daily consumption of food by rats, so that the average daily dose of the drug was ∼20 mg/kg. Blood glucose and body weight were measured in all the rats at the end of the pre-treatment period under 2% isoflurane anaesthesia.

### Ischaemia/Reperfusion model

Rats were anaesthetised with isoflurane (4% for induction and 2% for maintenance) and subjected to left anterior descending artery (LAD) ligation for 40 min, followed by 2hr of reperfusion. Blood pressure and ECG were recorded throughout the experiment, and body temperature maintained at 35.4 to 37.0°C. At the end of the reperfusion period, the animals were euthanised with an overdose of pentobarbital and the LAD was re-occluded. The heart was then perfused via the jugular vein with 5% Evan’s blue dye to delineate the area at risk (AAR). Infarct size (IS) was detected using 1% TTC staining.

### The effect of vagotomy and systemic non-selective muscarinic receptor blockade on cardioprotection established by SGLT2 inhibition

Both vagal nerves were exposed at the neck level prior to the experiment and sectioned simultaneously 10 min before myocardial ischaemia^26,27^ (**Fig. 1A**). In a separate group of animals, a non-selective muscarinic receptor blocker, atropine sulphate, was administered as an initial i.v. bolus of 2 mg/kg 10 min before myocardial ischaemia, followed by an infusion at a rate 1 mg/kg/h, as previously described^26^.

**Figure 1:**
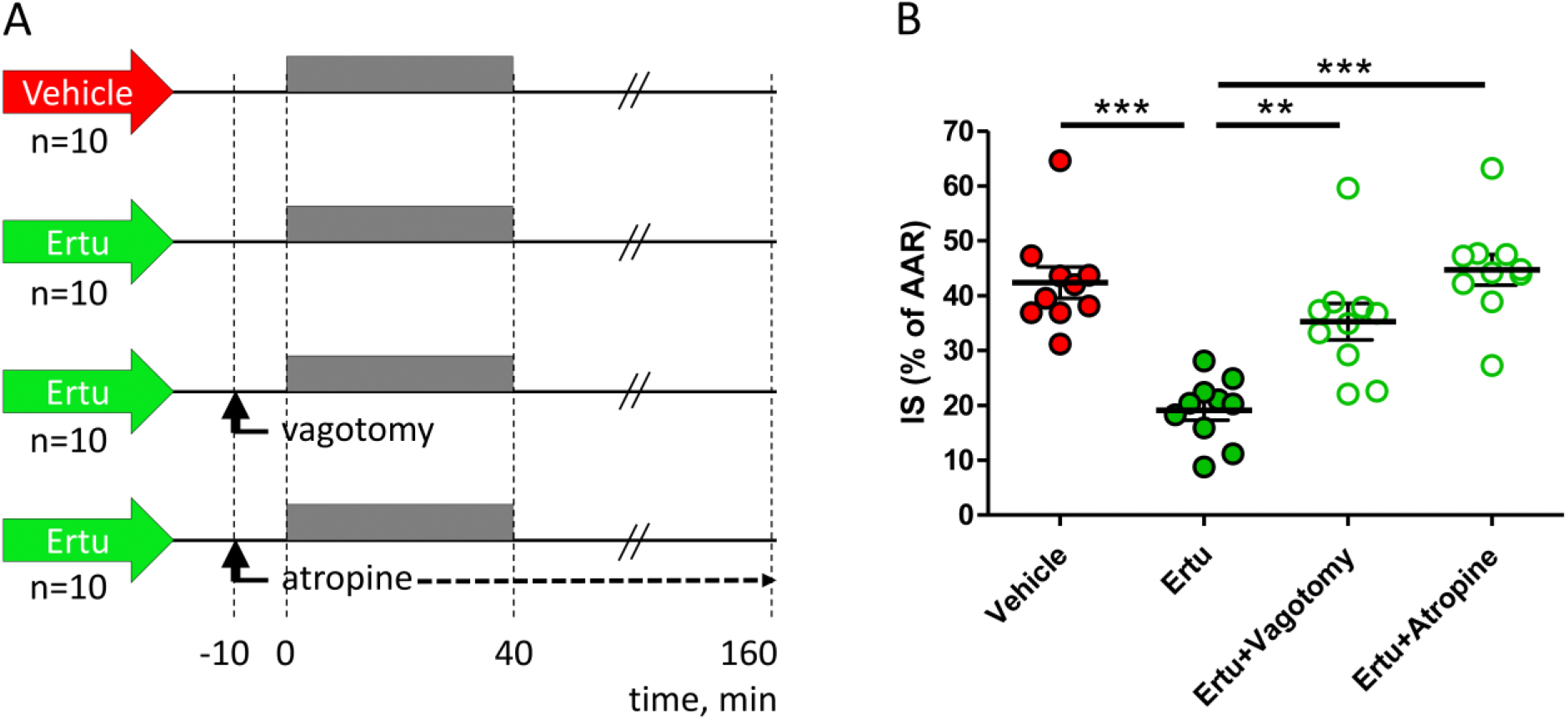
Pre-treatment with ertugliflozin (Ertu) reduces IS in SD rats subjected to 40 min myocardial ischaemia followed by 2hr reperfusion. This infarct-limiting effect of ertugliflozin was abolished by either bilateral cervical vagotomy or non-selective muscarinic blockade with atropine, or selective M3 muscarinic blockade 4-DAMP. IS was detected with TTC staining and measured by planimetry. The difference in IS between the groups was evaluated using 1 way ANOVA and Tukey post test. ** - P<0.01, *** - P<0.001.

### The effect of M3-muscarinic receptor blockade on cardioprotection established by SGLT2 inhibition

In a third group of animals, the selective M3-muscarinic receptor antagonist 1,1- Dimethyl-4-diphenyl acetoxypiperidinium iodide (4-DAMP) was administered as an i.v. bolus of 2 mg/kg 10 min before myocardial ischaemia (**Fig. 2A**).

**Figure 2:**
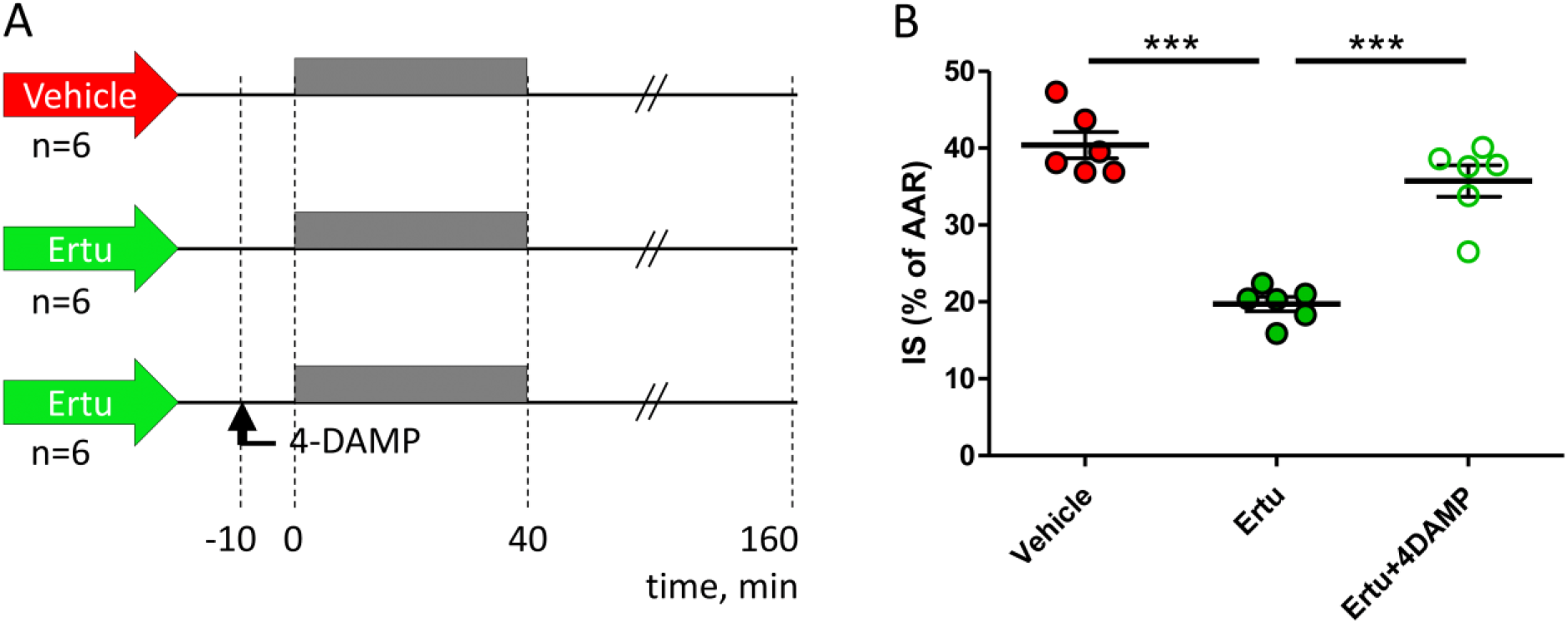
Pre-treatment with ertugliflozin (Ertu) reduces IS in SD rats subjected to 40min myocardial ischaemia followed by 2hr reperfusion. This infarct-limiting effect of ertugliflozin was abolished by selective M3 muscarinic blockade with 4-DAMP. IS was detected with TTC staining and measured planimetry. The difference in IS between the groups was evaluated using 1-way ANOVA and Tukey post test. *** - P<0.001.

### Randomisation and blinding

Rats were randomly allocated to the experimental groups. The experimenter was not blinded to the experimental groups, however IS was subsequently evaluated in heart slices while blinded to treatment group.

### Statistical analysis

Individual data points are shown as well as mean and standard error of the mean. Data was analyzed using 1-way ANOVA and Tukey post test. Significance is indicated by: ** - P<0.01, *** - P<0.001.

## Results

No difference in body weight was observed between the rats pre-treated with vehicle or ertugliflozin. Blood glucose in all the included animals at the end of the pre-treatment period varied in the normal range (7-10 g/mol) and was similar in the rats pre-treated with vehicle and ertugliflozin.

### The effect of vagotomy and systemic non-selective muscarinic receptor blockade on cardioprotection established by SGLT2 inhibition

Pre-treatment with ertugliflozin (Ertu) reduced IS by 55% (p<0.001 *vs*. vehicle) (**Fig. 1B**). Both bilateral vagotomy and atropine sulphate abolished the infarct-limiting effect of ertugliflozin (IS 35±10% and 44±8% respectively, P<0.01 and P<0.001 *vs*. Ertu; neither of the inhibited groups were significantly different from vehicle).

### The effect of M3-muscarinic receptor blockade on cardioprotection established by SGLT2 inhibition

In this series of experiments, pre-treatment with ertugliflozin (Ertu) reduced IS by 53% (P<0.001 *vs*. vehicle) (**Fig. 2B**). Selective M3-muscarinic receptor antagonist 4-DAMP abolished this infarct-limiting effect of ertugliflozin (IS 35±4%, P<0.001 *vs*. Ertu, P>0.05 *vs*. vehicle).

## Discussion

The key finding of this study is that activation of the parasympathetic nervous system is an important mechanism of the infarct-limiting effect of the SGLT2 inhibitor ertugliflozin. Cardioprotection established by ertugliflozin was found to be abolished both by vagotomy and systemic non-selective muscarinic receptor blockade with atropine. Importantly, this cardioprotective effect of ertugliflozin was also abolished using the selective M3 muscarinic receptor blocker, 4-DAMP. This may suggest that cardiac M3-receptor mediated mechanism is crucial for cardioprotection induced by SGLT2 inhibitors. We believe that this group of drugs may activate vagal efferent fibers innervating the ventricles, triggering the release of ACh in the myocardium, which protects ventricular cardiomyocytes via activation of M3 muscarinic receptors, which, apart from other functions, are known to mediate protection from ischaemia/reperfusion injury ^24,25^. Intriguingly, parasympathetic activation has previously been identified by our research group as crucial for the infarct-limiting effect of remote ischaemic conditioning^27,28^ and glucagon-like peptide-1 receptor agonist Exendin-4^26^. It cannot be excluded that this mechanism of cardioprotection is common for many other therapeutic interventions demonstrating reduced myocardial damage in the setting of acute coronary syndromes.

As yet, our data do not answer the question as to how SGLT2 inhibitors activate the vagus nerve. The main target organ for this group of drugs is the kidneys, which demonstrate the highest expression of SGLT2^29^. However, kidneys have no parasympathetic innervation, and are innervated with sympathetic fibers^30^. Intriguingly, a recent experimental study demonstrated that SGLT2 is abundantly expressed in the central nervous system (CNS) nuclei related to autonomic control^31^, although their role in the CNS is not clear. In mice treated with an SGLT2 inhibitor, an increase in neuronal activity was detected in a number of locations including the locus coeruleus, lateral parabrachial nucleus, the nucleus of the solitary tract (NTS) and the thalamic reticular nucleus. These nuclei relay and process afferent sensory signals to the neurons of the NTS and then activate interneurons to send the signals to other brainstem vagal nuclei such as the nucleus ambiguous and dorsal motor nucleus of vagus nerve (DVMN)^31^. As a result, this may lead to an increase in parasympathetic outflow to the heart^32,33^, which could promote cardioprotection^33,34,35^. Our research group have previously demonstrated that optogenetic stimulation of the DVMN vagal neurones reduces IS in a rat model of myocardial ischaemia/reperfusion^28^. Based on the above, we therefore hypothesize that activation of vagal pre-ganglionic neurons in the DVMN may be the crucial mechanism mediating the infarct-limiting effect of SGLT2 inhibitors in the heart.

Even though this hypothesis seems plausible, we cannot exclude the involvement of a more complex neuroendocrine loop into the mechanism of cardioprotection by SGLT2 inhibitors. Namely, SGLT2 inhibitors may stimulate vagal afferents in the intestine or other visceral organs under the diaphragm, and this may lead to the activation of vagal efferents via the mechanism of vago-vagal reflex^36^. In its turn, activation of subdiaphragmatic vagus efferents may enhance release of GLP-1^37,38^ and thereby launch the mechanism of cardioprotection similar to that underlying remote ischaemic conditioning^39,26^.

## Conclusion

The results obtained suggest that the infarct-limiting effect of SGLT2 inhibitor ertugliflozin is mediated via activation of the vagus nerve and M3-muscarinic receptors.

## Acknowledgement

The study was supported by a research grant from MSD.

